# STING suppresses migration of murine triple-negative breast cancer cells E0771 and 4T1 in vitro

**DOI:** 10.64898/2026.03.17.711042

**Authors:** Jason Xie, Neal Tandon, Yutong Li, Jean Zhao

## Abstract

Triple-negative breast cancer (TNBC) is the most aggressive subtype of breast cancer and lacks effective therapies. The stimulator of interferon genes (STING) has been shown to both suppress and promote migration in various cancer types, but its role in TNBC remains unclear. To investigate this, we established STING-overexpressing murine TNBC cell lines and assessed their migratory and proliferative behavior. STING overexpression significantly suppressed cell migration without affecting cell proliferation. Furthermore, STING overexpression in E0771 cells upregulated expression levels of *Itgb1* and *Itga6* significantly, but not *Icam1, Cxcl3, Itgb2, Lama5*, or *Rhoa*. These findings highlight the potential anti-migratory role of STING beyond immunomodulatory functions.

## Description

Triple-negative breast cancer (TNBC) is highly metastatic, and tumor-cell migration is a key determinant of disease progression and patient outcome (van den Ende et al., 2023). Among innate immune pathways, the stimulator of interferon genes (STING) has emerged as a key regulator of cell motility (Hu et al., 2023; Yang et al., 2023; Zhao et al., 2025). However, STING’s role in migration is context-dependent: it has been reported to either suppress or promote migration and invasion, dependent on the cancer type and experimental conditions (Zhou et al., 2023; Li et al., 2023; Zhao et al., 2025). Notably, STING expression is often downregulated in metastatic TNBC, suggesting a potential link between STING loss and enhanced migratory behavior.

Given these conflicting findings, we investigated the functional role of STING in TNBC cell migration. Using murine TNBC lines E0771 and 4T1, we established STING-overexpressing models.

We first confirmed successful STING overexpression. Western Blotting of whole-cell lysates showed elevated STING in STING-transduced cells compared to controls in both E0771 and 4T1 (Fig. 1A).

**Figure 1.**
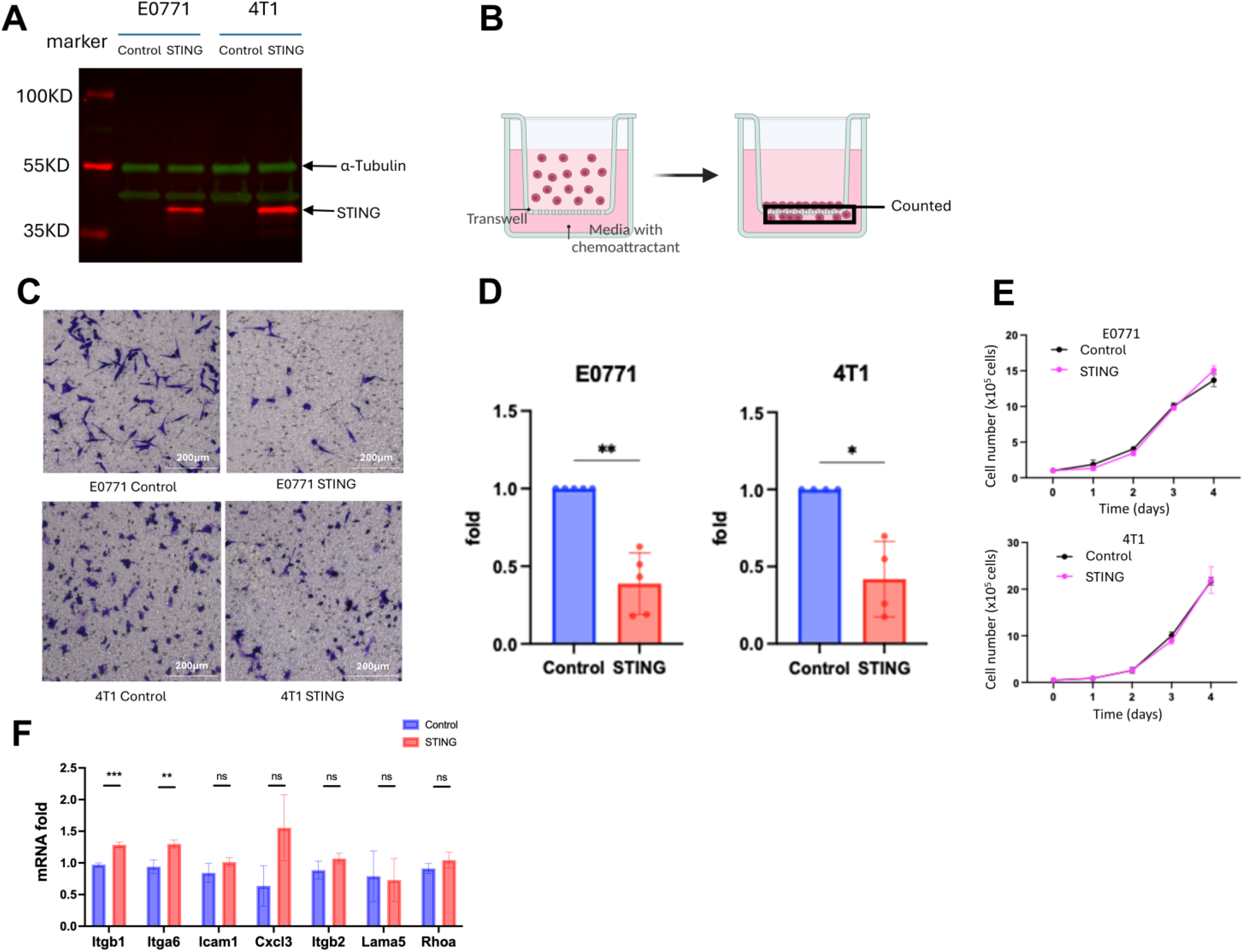
**(A) STING expression in STING-overexpressing and control TNBC cells**. α-Tubulin serves as a loading control. **(B) Transwell migration schematic**. **(C) Representative images from migration assays in STING-overexpressing and control TNBC cells (E0771 and 4T1)**. Scale bar = 200 µm. **(D) Quantification of cell migration in STING-overexpressing and control TNBC cells (E0771 and 4T1)**. Bars show mean fold change in migration with individual biological replicates overlaid and normalized to their matched control mean. E0771: n = 5, two-tailed paired Student’s t test, p = 0.0023. 4T1: n = 4, two-tailed paired Student’s t test, p = 0.0178. Data are presented as mean ± SD. **(E) Cell proliferation of STING-overexpressing and control TNBC cells (E0771 and 4T1)**. Growth curves over 4 days based on cell counts for the indicated cells. Data are presented as mean ± SEM. **(F) qRT-PCR analysis of gene expression in STING-overexpressing and control TNBC cells (E0771)**. *Itgb1* (n = 3, p < 0.001) and *Itga6* (n = 3, p < 0.01) show statistically significant increases in fold change relative to control in qRT-PCR. Data are presented as mean ± SD.

Next, we performed transwell migration assays (Fig. 1B) to assess the effect of STING on cell migration. STING-overexpressing cells exhibited significantly reduced migration compared with controls, with ∼61.2% reduction (n = 5, p = 0.0023) in E0771 cells and ∼58.0% reduction (n = 4, p = 0.0178) in 4T1 cells (Fig. 1C and D). These results demonstrate that tumor-intrinsic STING reduces cell migration in two distinct TNBC cell lines.

To address whether decreased transwell counts reflected differences in proliferation rather than motility, we measured cell growth over four days under the same culture conditions used for migration assays. Growth curves for STING-overexpressing and control populations were comparable in both E0771 and 4T1 (Fig. 1E). Therefore, reduced transwell migration was not due to altered cell proliferation.

To explore potential mechanisms, we quantified mRNA levels of a set of adhesion/motility-related genes by qRT-PCR. STING-overexpressing E0771 cells displayed significant upregulation of *Itgb1* (p < 0.001) and *Itga6* (p < 0.01), while the expression levels of *Icam1, Cxcl3, Itgb2, Lama5*, and *Rhoa* did not differ from controls (Fig. 1F). These results suggest that increased integrin-related transcripts are not associated with enhanced migration, implying potential post-transcriptional regulation or noncanonical mechanisms underlying STING-mediated migration suppression.

How STING modulates cell migration remains poorly understood. In this study, we found that STING overexpression selectively upregulated gene expression of *Itgb1* and *Itga6*, but not *Icam1, Cxcl3, Itgb2, Lama5, or Rhoa*. This complexity warrants further mechanistic investigation, including evaluation of downstream signaling pathways and protein-level changes. Moreover, as our study was conducted in vitro, in vivo animal models will be critical to validate the clinical relevance of these findings. Importantly, we assessed the effect of STING overexpression rather than activation of the cGAS-STING pathway, which may produce distinct biological effects. Future studies should compare the effects of STING overexpression versus pharmacologic or cyclic dinucleotide-induced activation in both immune and tumor compartments.

In conclusion, our data show that STING overexpression reduces cell migration in E0771 and 4T1 TNBC cells in vitro without affecting proliferation. In E0771 cells, STING overexpression was associated with upregulation of specific integrin-related genes, including *Itgb1* and *Itga6*. These findings support a tumor-intrinsic, immune-independent role for STING in regulating cancer cell motility. Further work should explore mechanistic studies and assess the impact of STING activation in vivo to establish its therapeutic potential in metastatic TNBC.

## Methods

### Cell lines and culture

Murine TNBC cell lines E0771 and 4T1 stably transduced to overexpress murine STING or empty vector control were generated in the Zhao Lab, Dana-Farber Cancer Institute. All cells were cultured in RPMI-1640 medium supplemented with 10% fetal bovine serum (FBS) and maintained at 37°C with 5% CO_2_. Cells were passaged at 80-90% confluency using 0.05% trypsin-EDTA and counted with an automated cell counter using trypan blue exclusion to assess viability.

### Transwell migration assay

Transwell inserts with 8.0-µm porous polycarbonate membranes (6.5 mm diameter, tissue culture-treated, Corning #3422) were placed in a 24-well plate. The lower chamber was filled with 700 µL of complete medium containing 10% FBS as a chemoattractant. Cells were prepared following the passaging protocol, centrifuged at 500 × g for five minutes, and resuspended in serum-free medium. A total of 100 µL of the cell suspension was seeded onto the upper chamber of each transwell insert. After 24 hours of incubation (E0771) or 6 hours (4T1), non-migrated cells on the upper surface of the membrane were removed with a cotton-tipped applicator. Migrated cells on the bottom side were fixed with 10% formalin for 10 minutes and stained with crystal violet for 2 minutes. Excess stain was removed by washing with water, and transwells were allowed to dry before imaging using a microscope. Biological replicates (n = 5 for E0771, n = 4 for 4T1) were performed on different days.

### Western Blot analysis

Cells were harvested and lysed in 1X loading buffer and then denatured at 100°C for five minutes. Samples were loaded onto Mini-PROTEAN TGX 4-20% polyacrylamide gels (Bio-Rad Laboratories) alongside a protein ladder (PageRuler Plus, Thermo Fisher Scientific) and resolved via SDS-PAGE at 75 V for 10 minutes, followed by 150 V for 60 minutes. Proteins were transferred to 0.45-µm nitrocellulose membranes (Thermo Fisher Scientific) at 100 V for 60 minutes with cooling. Following transfer, membranes were blocked in a 1:1 mixture of Odyssey blocking buffer (LI-COR Biosciences) and PBS for 30 minutes at room temperature, then incubated overnight at 4°C with primary antibodies: anti-STING (Cell Signaling Technology #13647, 1:1,000) and anti-α-tubulin (Sigma-Aldrich T9026, 1:5,000). After three washes with PBS-T, membranes were incubated with fluorescent secondary antibodies (goat anti-mouse IgG-DyLight 800 and goat anti-rabbit IgG-DyLight 680) (1:7,500 each) for 30 minutes at room temperature, protected from light. Following three additional PBS-T washes and a final PBS rinse, fluorescence was detected using LI-COR Odyssey imaging system.

### Cell proliferation assay

Cells were seeded in 6-well plates at a density of 1 × 105 cells/well (E0771) or 5 × 10^4^ cells/well (4T1). Cells were trypsinized and counted daily for 4 days using 0.4% trypan blue staining and a Countess II FL automated cell counter (Thermo Fisher Scientific). Experiments were performed in triplicate for E0771 (n = 3) and quadruplicate for 4T1 (n = 4).

### Quantitative reverse transcription polymerase chain reaction (qRT-PCR)

Total RNA was extracted from cell lysates using Trizol reagent (Thermo Fisher Scientific) followed by RNeasy mini kit (Qiagen) and quantified using a NanoDrop spectrophotometer. Reverse transcription was performed to generate complementary DNA (cDNA) using Superscript III First Strand Synthesis System (Thermo Fisher Scientific). Quantitative PCR was performed in triplicate using PowerUp SYBR Master Mix (Thermo Fisher Scientific) and gene-specific primers (Integrated DNA Technologies) for *Itgb1, Itgb2, Itga6, Icam1, Cxcl3, Lama5*, and *Rhoa*. Reactions were run on a real-time PCR system under optimized cycling conditions. Threshold cycle (Ct) values were obtained and normalized to housekeeping gene *Gapdh*, with relative expression calculated using the ΔΔCt method (Livak & Schmittgen, 2001).

### Automated segmentation

Stained transwell images were segmented using Cellpose 2.0 graphical user interface human-in-the-loop (HITL) workflow (Pachitariu & Stringer, 2022). We conducted a density screen in ImageJ (Schindelin et al., 2012) to select representative fields, and quantitative counts were derived from Cellpose masks following manual quality control. A curated training set was built from transwell imaging, and the generalist Cellpose model was fine-tuned across multiple iterative rounds of training (Stringer et al., 2021; Pachitariu & Stringer, 2022). The following parameters were used: 1200 epochs and learning rate = 9.8 × 10^-4^.

### Analysis

Migration data are presented as fold change relative to control (set to 1.0). qRT-PCR data are presented as fold change relative to control using the ΔΔCt method. Statistical significance was assessed using two-tailed paired Student’s t test in GraphPad Prism. p < 0.05 was considered significant. Plots were generated using GraphPad Prism.

## Reagents

qRT-PCR primers used in this study are listed below.

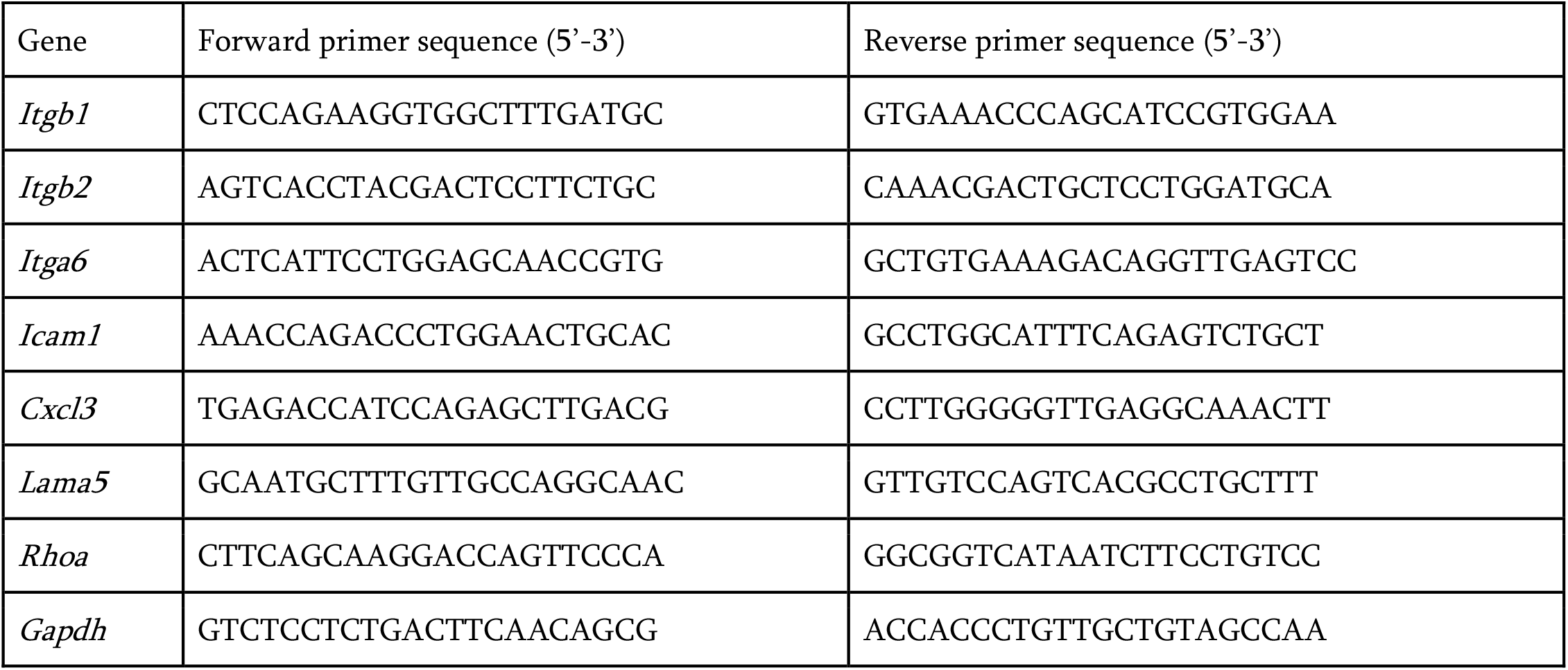

## Acknowledgements

We thank Kenneth Macie (Dover-Sherborn High School) for guidance throughout this process.

## Funding

This work was supported by the Dana-Farber Cancer Institute.

## Author Contributions

Jason Xie: Conceptualization, Investigation, Formal Analysis, Visualization, Writing – original draft

Neal Tandon: Conceptualization, Data Curation, Formal Analysis, Visualization

Yutong Li: Methodology, Supervision, Validation, Writing – review & editing

Jean Zhao: Conceptualization, Supervision, Project administration, Resources, Writing – review & editing

## Notes

### Competing Interest Statement

The authors have declared no competing interest.

### Summary of Updates

This version revises the Abstract, Figure 1 legend, and concluding text to clarify the scope of the gene expression findings and improve consistency with the data shown. Minor wording, methods, and formatting edits were also made throughout. These changes do not affect the main results or conclusions of the manuscript.

